# bbsBayes: An R Package for Hierarchical Bayesian Analysis of North American Breeding Bird Survey Data

**DOI:** 10.1101/2020.05.27.118901

**Authors:** Brandon P.M. Edwards, Adam C. Smith

## Abstract

The North American Breeding Bird Survey (BBS) is the primary ecological monitoring program used to assess the population, status, and trend of North American birds. As such, accessible analysis of BBS data is crucial to wildlife conservation/management and ecological science in North America. The R package *bbsBayes* was developed as a wrapper for the analysis of BBS data using hierarchical Bayesian models, including the models currently used by the Canadian Wildlife Service and the United States Geological Survey. The goal of *bbsBayes* is to provide an accessible package for anyone in the conservation community to estimate population trajectories (time-series) and trends (rates of change) for any of the 400+ bird species monitored by the BBS, and to allow more advance users to easily access the data and model-templates necessary to customize an analysis for their research.

## 1.1 Introduction

The North American Breeding Bird Survey (BBS) is a long-running, large-scale data collection program for monitoring breeding bird populations of North America [10]. The BBS was started in 1966 and now provides a long-term database of counts of over 400 species of birds across more than 5000 BBS routes, in Canada and the United States, and a few in Mexico [6, 24]. BBS data are collected annually by hundreds of volunteers, who survey their assigned routes following a rigorously standardized protocol. Each BBS route follows a randomly selected, secondary road and consists of 50 stops, separated by approximately 800 m, where the observer conducts a 3-minute point count, counting all the birds they can see and birds they can hear within 400 m. The data collected by the BBS have been the primary source of data in more than 700 scientific publications to date [28].

Estimates of population trends (rates of change in population size) and the associated trajectories (series of annual estimates of relative abundance) are derived from the BBS data annually by federal government agencies in the U.S.A.[24] and Canada [27]. These trends and trajectory estimates are used by government, academic, and public organizations to monitor bird populations, set priorities for conservation, identify species at risk, and explore fundamental ecological relationships [6]. For example, BBS status and trend estimates were a major component of the study led by Rosenburg et al (2019) that showed a decrease of 3 billion birds since 1970 [20]. BBS status and trend estimates also provide the large majority of the information used in State of Birds reporting in the U.S.A. and Canada. [13, 14] And, in Canada, estimates of trends are used by the Committee on the Status of Endangered Wildlife in Canada (COSEWIC) to inform decisions as to whether species should be added or removed from the list of species at risk [27]. The models and modelling frameworks used to produce estimates from the BBS are continually evolving [22, 23, 27, 26]. Currently, these trend and trajectory estimates are generated using a hierarchical Bayesian, over-dispersed Poisson regression model that accounts for variation among and within observers, routes, and geo-political strata [23, 27].

Generally, the raw BBS data and the trend and trajectory estimates have been made easily available through the CWS and USGS websites [15]; however, the detailed model structure, and the analytical tools necessary to produce these estimates are not nearly as accessible.

The United States Geological Survey (USGS) and Canadian Wildlife Service (CWS) jointly manage the BBS data and publish the entire data set each year in an online format that’s easily accessible to users [15]. However although the data are available and the models have been described in the literature, there does not yet exist a streamlined method for a researcher to download the entire data set and replicate the estimates of trends and trajectories, or to apply a customized hierarchical Bayesian model for a species of interest. Additionally, the process of properly subsetting and preparing the raw BBS data for use with JAGS models (for example, adding zero-counts) may be a daunting task to a researcher who may not have much experience using Bayesian models or exploring the BBS data.

The R package *bbsBayes* was developed to address these issues: to provide an easy-to-use R package that contains standard BBS models and analysis scripts that can be used by scientists interested in hierarchical Bayesian BBS data analysis. In this paper, we detail the development of the package itself, as well as detail the data prepping, modelling, and data analysis functionality that can be performed with *bbsBayes*. Using the functionality described, we provide a worked example of a full analysis run on Wood Thrush (*Hylocichla mustelina*), a near-threatened and declining passerine bird that breeds in North America [7].

### 1.2 Implementation and architecture

*bbsBayes* seeks to mirror the typical workflow of BBS data analysis for a given species. As such, the package provides functionality for 4 main processes:

1. Data import
2. Data preparation
3. Markov chain Monte Carlo (MCMC) simulation
4. Model summary

The user has access to functionality under all four processes to analyse data as needed for their research. The following subsections detail the implementation of each of these processes and describe the functionality available to the user from each process. We will also give a brief discussion on how to modify the models to add covariates, change priors, etc.

#### 1.2.1 Data Import

When the package is first installed, the user must download the data from the USGS, which can be accomplished within the *bbsBayes* package using the `fetch_bbs_data()` function. When called, this function downloads raw BBS data from the USGS website via File Transfer Protocol (FTP) and saves it to an application directory on disk. Note that this function must be run upon the user’s first use of *bbsBayes*, and it is recommended to rerun this function each year as USGS publishes updated data sets.

To bring the raw, unstratified data into R, the function `load_bbs_data ()` can be used. Three data frames are created from this function – `route`, `bird`, and `species` – and are returned as a large list. Since BBS data is well documented by the USGS [15], we will not detail every variable for each data frame. Instead, we will highlight key variables that are used in data preparation and simulation.

##### 1.2.1.1 Route Data

The `route` data frame provides a variety of information for each route run by year. Each entry in `route` corresponds to route information per year, so route *x* run in year *y* will have its own row unique from route *x* run in year *y* + 1. Each route will have corresponding locational data with it, such as state/province, country, latitude/longitude, and Bird Conservation Region (BCR, [2]) number.

Each route is designated a unique name, usually referring to the general location of the route. In addition, the routes are assigned numbers which are unique only to the state/province it is in; for example, route *x* in Alabama could be assigned as route 1, and route *z* in Ontario could also be assigned route 1. By combining route number, state, and year, we can get a unique identifier for each route by state run in each year.

Each entry in `route` also contains information about the weather observed on the route. Currently, these data are not used in analyses, but are at least available for use by other scientists.

##### 1.2.1.2 Bird Data

The `bird` data frame provide species counts by route. Each row of the `bird` data frame corresponds to the number of individuals observed of a given species on a given route, in a given year. For example, the number of mallards observed on route *x* in year *y* will be its own row separate from the number of mallards observed on route *x* in year *y* + 1. Similarly, if 50 species were observed on route *x* in year *y*, then 50 separate rows will exist with count data for each species on route *x* in year *y*. Note: this data frame does not include zero counts; if a species is not observed on a given route, in a given year, there will be no record of it in this file, even if that species has been observed on that route in other years.

The `route` variable in the bird data frame corresponds to the numerical representation of the route as described in 1.2.1.1. Additionally, each entry in bird is assigned general locational information, such as state/province, country, and Bird Conservation Region (BCR) number. Indeed, the combination of route number, state, and year can be used to cross reference species count data to the route data contained in the `route` data frame.

##### 1.2.1.3 Species Data

The `species` data frame simply provides a list of all North American bird species in their English, French, and Spanish names, their taxonomic classifications, and a unique numeric code assigned by the American Ornithological Union (AOU; now the American Ornithological Society, AOS).

#### 1.2.2 Data Preparation

Data preparation functions are provided for functionality to prepare raw BBS data into data used as input for JAGS modelling. As such, data preparation is heavily dependent on what analyses the user is looking to perform.

##### 1.2.2.1 Stratification

Stratification of BBS data is used to provide regional estimates for different geographical areas of North America. The function `stratify()` is used to create an intermediate `bird` and `route` data frame that contain references to the geographic stratum-allocation for each data point. There are two ways to use the function. In the first way, the user can load raw BBS data into the R session using the `load_bbs_data()` function and save it to a variable. This variable would then be passed into the `stratify()` function with the `bbs_data` argument. This approach would be most useful if an advanced user wished to make some modification to the dataset to suite a customized model or a custom subset of the data.

However, since `fetch_bbs_data()` saves raw BBS data directly to the user’s disk, functions from *bbsBayes* can therefore access this data. Indeed, the less memory-intensive and recommended way to use the `stratify()` function is to simply not provide any input data to the function; the function will access the raw data from the user’s disk, stratify the data, and return the stratified data as a large list.

To specify how the data should be stratified, the user provides a string to the by argument, which can be given 5 possible options:

1. “latlong” – Stratify by degree blocks (i.e. 1 degree of latitude by 1 degree of longitude, the sampling strata with which the BBS routes are defined)
2. “state” – political region only (i.e. by province, state, or territory)
3. “bcr” – Bird Conservation Region (BCR) only
4. “bbs_usgs” – Intersection of political region and BCR (USGS method)
5. “bbs_cws” – Intersection of political region and BCR (CWS method)

Figure 1 illustrates each of these stratification options on maps of North America.

**Figure 1:**
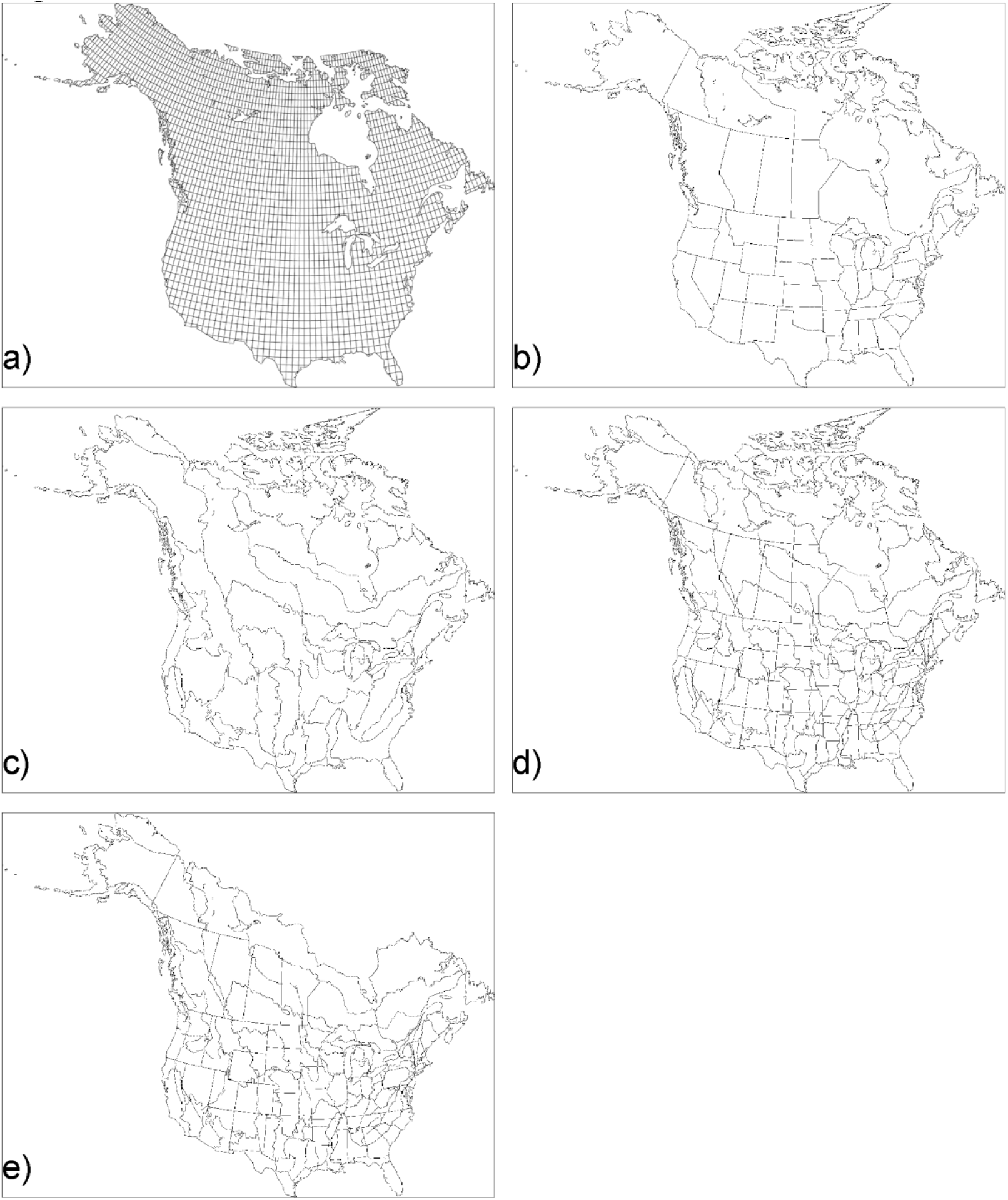
Maps of the 5 stratification options offered by *bbsBayes*. A user can specify to stratify by degree blocks (a), state (b), Bird Conservation Region (BCR; c), BCR X state (USGS method; d), or BCR X state (CWS method; e).

In current hierarchical Bayesian analyses of BBS data, option 4 (“bbs_usgs”) or option 5 (“bbs_cws”) are typically used. The USGS and CWS methods of these intersections vary slightly. In both, the strata are defined by the intersection of political borders (i.e. state, provincial, and territorial borders) and BCR borders. For example, for a route located in Toronto, Ontario, Canada, the political boundary is specified by the province of Ontario, and the BCR boundary is specified by BCR 13: Lower Great Lakes/St. Lawrence Plain, so all counts conducted in this intersection would be assigned to the stratum “Ontario-BCR13”. The CWS, however, makes 2 modifications to this method: 1) The provinces of Prince Edward Island and Nova Scotia are treated as a single province, because PEI is so small that it only contains 4 BBS routes and therefore often fails to meet the minimum data criteria; 2) The entirety of BCR 7: Taiga Shield and Hudson Plains is treated as one stratum.

Each entry of the `route` data frame now has a reference to the stratum that it is contained in. Each of the `bird` and `route` data frame gain a unique identifier for each entry that is a combination of the state, route, and year (as mentioned in 1.2.1). If the user selected to stratify using the CWS method of BCR and state intersection, both the `route` and `bird` data frames will be modified to reflect the combined provinces of PEI and Nova Scotia and the combined routes for BCR 7.

The `stratify()` function returns a large list that contains modified versions of the `bird` and `route` data frame that contain references to their strata.

##### 1.2.2.2 JAGS Preparation

The function `prepare_jags_data()` creates the species-specific data used for fitting the JAGS model. The models supplied by *bbsBayes* can only be used for one species at a time, so this function must be called for each species to be modelled. This function provides the user with options to select the species to model (by setting the `species` argument), as well as data exclusion criteria and options for model-specific required data.

`prepare_jags_data()` uses the stratified data set created by `stratify()`. The function starts simply by subsetting the `bird` data frame, keeping only observations of the specified species. This subset of data is then merged with the `route` data frame. As mentioned in 1.2.1, the `bird` data frame and `route` data frame both contain references to the state/province and BCR the data were recorded in (i.e. what state/province and BCR the route is in), the year for each route run, and a numerical representation of the route itself. Therefore, these two data frames can be merged by these 4 variables.

The merging of these two data frames serves two purposes. For one, since the strata information is contained in the `route` data frame (among other qualitative data about the route), merging these two data frames now allows for a strata reference for the species of interest, and allows a reference for qualitative route data as possible covariates for the species count. Additionally, merging these two data frames adds the zero-counts to the data. Because the list of species not observed on any given route *x* year combination is extremely long, zero-counts are not explicitly stored in the BBS data set: an entry for species s on route *x* in year *y* will appear in the `bird` data frame only if it was actually observed. This creates the case where if species s was only observed for 10 out of the 50 years that route *x* has been run, there are implied yet missing data points that shows 0 count for the remaining 40 years. With this merge, we ensure that all the years a given route was run appear in the merged data frame, even if the species was not observed in a particular year.

`prepare_jags_data()` provides a number of arguments for the user to exclude data from the analysis that do not meet a threshold. These are:

- `min_year`: Set the minimum year to keep in the analysis
- `max_year`: Set the maximum year to keep in the analysis
- `min_n_routes`: specify the minimum routes per strata where the species has been observed
- `min_max_route_years`: specify the minimum number of years with nonzero observations of the species on at least 1 route
- `min_mean_route_years`: specify the minimum average of years per route with the species observed
- `strata_rem`: specify which stratum or strata to withhold from the analysis

The authors of *bbsBayes* have set defaults for these variables based on what is done for analyses performed by CWS.

*bbsBayes* incorporates four Bayesian models into the package. When the user calls `prepare_jags_data()`, they will need to specify for which model the data will be prepared using the `model` argument. As of this version (additional models will be added in the future), there are 4 model options, reflecting 4 different approaches to modelling the temporal patterns of population change (Table 1). For each model, the user can also specify whether the extra-Poisson error distribution should be modelled as a normal distribution or a heavy-tailed t-distribution, the latter of which is accomplished by setting `heavy_tailed = TRUE`.

**Table 1:**
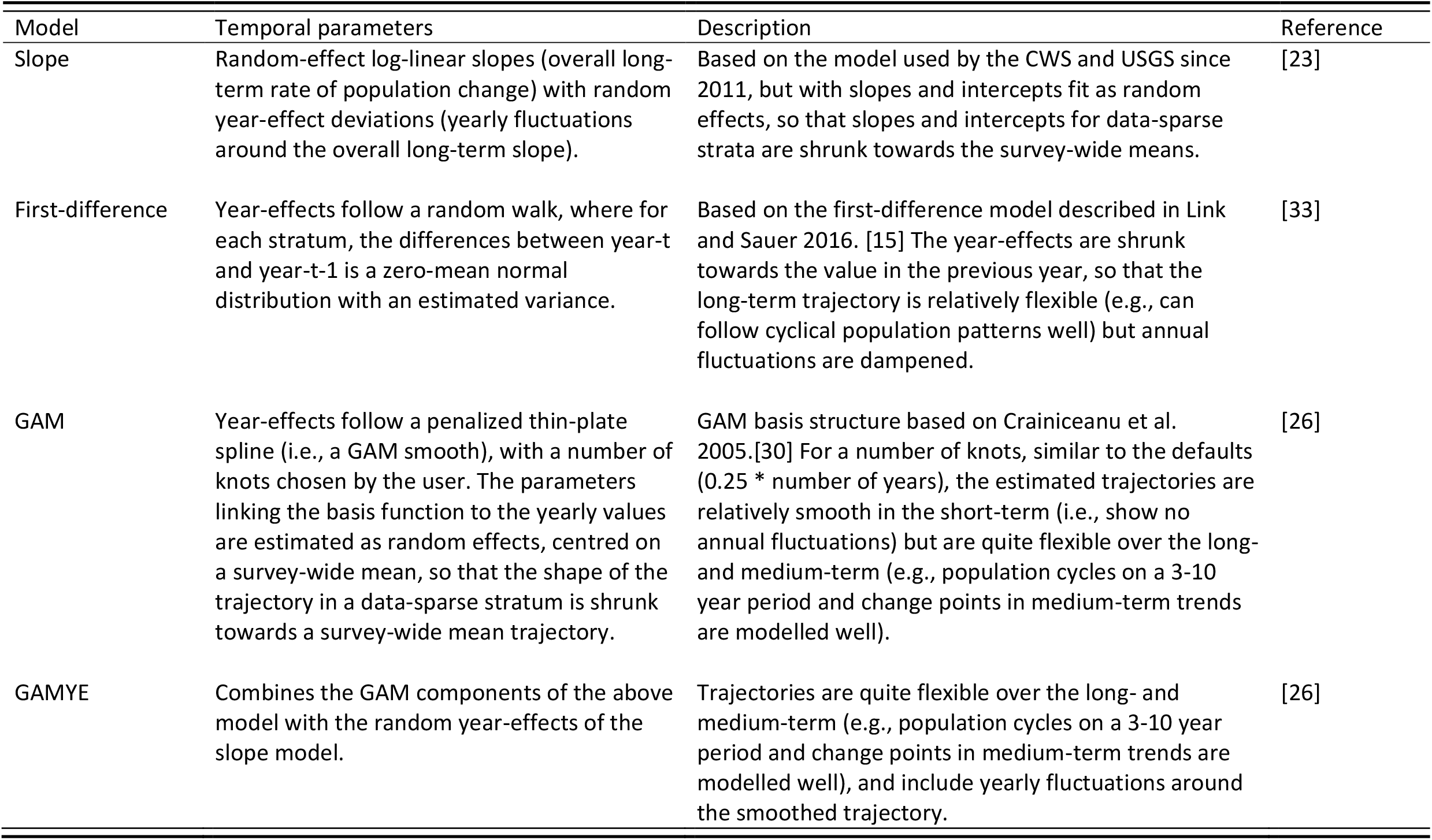
Comparison of the 4 models provided by *bbsBayes*.

The generalized additive model (GAM) and GAM year effect (GAMYE) require a basis function with *n* number of knots [4]. The user can specify the number of knots to be used with the `n_knots` argument, but `prepare_jags_data()` will default to 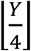, rounded to the nearest integer, where *Y* is the total number of years.

In all four models, the data list returned by `prepare_jags_data()` will contain the number of counts, number of strata used, the area-weight of each stratum, minimum and maximum years, the count data per route per year, the stratum for each count, the observer-route combination for each count, the year for each count, the number of observers for each count, and an indicator variable of whether it was the observer’s first year of counting. If the user chooses the slope model, a fixed year, which is simply the median of all the years, is added into the list. If the user chooses the GAM or GAMYE models, the basis function matrix will be returned in this list.

#### 1.2.3 MCMC Simulation

MCMC simulation is accomplished using functionality from the *jagsUI* package [11]. The function `run_model()` acts as a wrapper for creating, adapting, and sampling from a JAGS model.

For simplicity, `run_model()` provides a variety of defaults: the only required argument to the function is the list of data produced by `prepare_jags_data()`. The model to be run is extracted from that list. `run_model()`, by default, will run a model with 3 chains, 20000 burn-in steps per chain, 10000 sampling iterations per chain. The model will thin each chain by 10 steps and will save a total of 2000 steps per model run. No initialization values are used. The function defaults to only tracking the derived parameter *n* (i.e., the trajectories of stratum-level annual indices of abundance), which is necessary for later trend analysis. *bbsBayes* offers a total of four possible ways to calculate this annual index of abundance:

1. `n:` Index of abundance as used in [27]. If the GAMYE model is chosen, this index includes the added random year effects.
2. `n2:` Index of abundance as used in [23]. If the GAMYE model is chosen, this index includes the added random year effects.
3. `n3:` Same index as n; however, does not include the random year effects if the GAMYE model is chosen (i.e., this is just the smooth component of the trajectory).
4. `n4:` Same index as n2; however, does not include the random year effects if the GAMYE model is chosen (i.e., this is just the smooth component of the trajectory).

By tracking both `n` and `n3` (or `n2` and `n4`), the user can decompose the GAMYE into a trajectory with random year effects and a trajectory that is just the GAM smooth [12, 26]

The user has the option to provide their own values for any of the JAGS parameters mentioned above. Advanced users may specify a character vector of JAGS modules to load before analysis. By default, no extra modules are loaded (other than “basemod” and “bugs”). To force “glm” or other modules to load, use `modules = “glm”`. The authors note, however, that including the “glm” module may cause problems with the BBS data.

The MCMC simulation is, by far, the most time consuming and computationally expensive process in the analysis of BBS data, with model runs of data rich species (such as Barn Swallow, Mourning Dove, or American Robin), and particularly with more complicated models (e.g., GAMYE), taking up to 24 hours or more to complete. If the user has more than 1 processor core available, the user may specify the argument `parallel = TRUE` to run one chain per processor to cut down significantly on this processing time. Users should pay close attention to the system requirements outlined in section 2.3.

#### 1.2.4 Model Summary

A number of model summary and visualization tools are available from *bbsBayes* to allow the user to generate meaningful metrics given the simulated posterior generated by `run_model()`.

##### 1.2.4.1 Convergence

In this package, we use Gelman and Rubin’s 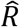(“R-hat”) statistic, also known as potential scale reduction factor (PSRF), to assess convergence 8]. R-hat is a ratio of the posterior variance estimates for the pooled traces of the parameter and the within-chain variance. When converged, both variance estimates will be equal, giving an R-hat value of 1. Values of R-hat greater than 1 imply that some chains may not have converged, and more sampling may be necessary [8].

`The run_model()` function will send a warning if 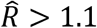 for any of the monitored parameters. From here, the user should decide as to how to proceed with model summary. To possibly improve convergence, one may consider taking more samples from the posterior; *bbsBayes* provides a `get_final_values()` function that will return the final values of a JAGS model which can be used as initial values to a new call to `run_model()`, avoiding the need to wait for an additional burn-in period (see 1.3 Worked Example: Wood Thrush).

However, the seriousness of a convergence failure is something the user must interpret for themselves. In some cases, some parameters of the model may not be separately estimable, but if there is no direct inference drawn from those separate parameters, their convergence may not be necessary. If all or majority of the n parameters have converged (e.g., the user receives a warning message for other monitored parameters), then inference on population trajectories and trends from the model are reliable.

##### 1.2.4.2 Generating Indices and Trends

Given the JAGS model output by `run_model()` and the prepared data for JAGS output by `prepare_jags_data()`, the function `generate_indices()` will output a data frame of strata-weighted indices as well as a vector of quantiles sampled from the posterior (to be used for plotting credible intervals about a trajectory). By default, the function will output indices for the continent (survey-wide) and for individual strata. However, the user can set the `regions` argument to output indices for composite regions such as countries, provinces and states, Bird Conservation Regions, etc.

Once the annual indices of abundance are calculated, they can be used to generate population trends. In *bbsBayes*, and in analyses performed by CWS and USGS, the trends are calculated as the geometric mean rates of change (% / year) between two points in time. This calculation is performed using the `generate_trends()` function, which can simply take the indices generated previously as input. The user can also specify a minimum year and maximum year for which to generate trends (by setting the `Min_year` and `Max_year` argument, respectively). The function will return a data frame with 1 row for each unit of the region types that were requested in the `generate_indices()` call (i.e., 1 row per stratum, 1 for continental, 1 per any other region selected). Additionally, the data frame contains other information related to each trend, including the start and end year of the trend, lists of included strata, total number of routes used, among others.

`generate_trends()` also allows for an alternative trend calculation. As mentioned, the default trend calculation is to calculate the geometric mean annual change in population between the start and end years. However, by setting the argument `slope = TRUE`, the function will also fit a log-linear slope to the series of all annual indices between the two end points. For some models that contain strong annual fluctuations for which no decomposition is possible (for example, the first difference model), the slope trend may be a better measure of average population change, as it integrates the pattern of change between the two end points.

Finally, `generate_trends()` provides functionality for the user to make statements about percent change and the probability of change for a given species. This is accomplished by setting the `prob_decrease and/or prob_increase` arguments with a vector of probabilities (i.e., a vector of numbers between 0 and 100, inclusive). Then, the function will output (for each row in the data frame) the probability of that change. For example, if the user set `prob_increase = c(0,100)`, then the data frame will contain columns related to the probability of the species increasing (that is, a > 0% increase over the time period) and the probability of the species increasing more than 2-fold (that is, a > 100% increase over the time period). Section 1.3 will cover this in more detail.

##### 1.2.4.3 Model Visualizations

With any modelling of large data sets, and especially when talking about an increasing or decreasing population of an animal species in its habitat, it is crucial to have clear and easy-to-interpret visualizations of what the model produces for indices and trends.

*bbsBayes* comes with a variety of functions to visualize the indices and trends produced by the model. The `plot_indices()` function is used to create a time series of the indices for each region specified in the initial call to `generate_indices()`. That is, given the data frame of indices, `plot_indices()` will return a list of plots (created with ggplot2 [30]), one per stratum, one continent-wide, and one for each additional region specified. The user has a variety of additional visual options for the trajectory plots: they can set `add_observed_means = TRUE` to overlay the observed mean counts for each year, they can set `add_number_routes = TRUE` to superimpose a dotplot of the number of BBS routes included for each year, they can change the minimum and maximum year to plot by setting the `min_year` and/or `max_year` arguments, or they can change the sizes of axes, title, text, etc.

These trajectories can also be visualized on a geofacet plot using the `geofacet_plot()` function, which is a graphic that plots the state/province level population trajectories in facets arranged approximately in a geographical arrangement [9]. Again, this function simply takes in the indices generated by `generate_indices()`. This plotting can only be accomplished if the data was stratified by one of “state”, “bbs_cws”, or “bbs_usgs”.

Finally, the stratum-specific trends created by `generate_trends()` can be mapped out using the `generate_map()` function which creates a stratified continental heat map.

#### 1.2.5 Modifying Models

Often with Bayesian models, new research is published that lead to more informative or useful priors on estimated parameters. Or, researchers may be interested in parameterizing an already-existing model differently to experiment with model selection. It therefore makes sense that the models supplied with *bbsBayes* should be reasonably simple to customize, and that the customized model files can easily be used with the rest of the functions of *bbsBayes*.

Indeed, the function `model_to_file()` allows the user to save the model files supplied by *bbsBayes* as plain text files. Suppose we wanted to save the log-mean slope model file to disk. We could use this function setting the arguments `model = “slope”` and `filename = “slope_model.txt”`, which will save the model file to the current working directory as a file called “slope_model.txt”. Suppose some part of that file was modified to our liking (e.g. changing priors) and we renamed the file to “slope_model_modified.txt”. We could then run our model by setting the `model_file_path argument in run_model() as model_file_path = “slope_model_modified.txt”`.

### 1.3 Worked Example: Wood Thrush

We will now provide an example of using *bbsBayes* to generate a meaningful quantitative analysis of Wood Thrush trends over the time period of the BBS data collection. The Wood Thrush is a medium-sized neotropical migrant that occurs in eastern North America during its breeding season. Since the late 1970s, habitat destruction in both wintering and breeding grounds have contributed to a severe loss of abundance in this species [7]. BBS data on Wood Thrush is reasonably data rich, so it makes it a good species to model and pick up these trends over time. The script for this example, as well as the prepared data generated from `prepare_jags_data()` and the model output generated from `run_model()` is available at https://github.com/BrandonEdwards/bbsBayes-Wood-Thrush-Worked-Example.

#### 1.3.1 Data Fetching, Preparation, and Modelling

We start with calling the *bbsBayes* library, then making a call to download the BBS data from the USGS FTP site. As mentioned, this step is required for the first use of *bbsBayes*, and should only be used once each year as USGS releases updated data sets.

~~~
# Use bbsBayes library
library(bbsBayes)
~~~

~~~
# Download the data (requires internet connection)
fetch_bbs_data()
~~~

When `fetch_bbs_data()` is called, the user must agree to the terms and conditions of the data usage by typing “yes” in the console. Otherwise, the data will not be downloaded. Once downloaded, however, the BBS data is saved on disk to a package-specific directory that can be accessed by functions of *bbsBayes*.

Now that we have the raw data downloaded to disk, we can bring the data into R and stratify it. The function `stratify()` will accomplish both these tasks. For this worked example, we choose to stratify by the state X BCR intersections (CWS method), thus our argument for by will be the string “bbs_cws”. We recommend assigning the string “bbs_cws” (or which ever stratification the user is choosing) to a variable as it will eventually be used later in the analysis.

~~~
# Stratify the data
s <- “bbs_cws”
stratified_data <- stratify(by = s)
~~~

The variable `stratified_data` contains the stratified data which can now be prepared for use with JAGS. In this example, we will model Wood Thrush counts using the hierarchical Bayesian generalized additive model with year effects, requiring the string “gamye” to be passed as an argument. Additionally, we must now specify which species to subset and prepare for JAGS.

~~~
# Prep the data for JAGS
jags_data <- prepare_jags_data(strat_data = bbs_strat,
                          species_to_run = “Wood Thrush”,
                          model = “gamye”)
~~~

We are now ready to run the JAGS model. As mentioned, this will typically end up being the most time- and memory-consuming process in an analysis, so we recommend that the user pay close attention to 2.3 Additional system requirements before running any JAGS models. As mentioned, the prepared data and model output for this worked example have been made available at https://github.com/BrandonEdwards/bbsBayes-Wood-Thrush-Worked-Example, for readers that may want to skip over this step and try out the model summary and visualization features.

We could simply run the following code to run a model with default settings:

~~~
jags_mod <- run_model(jags_data = jags_data)
~~~

However, for this example, we will modify a number of default settings for the JAGS model. Here, we will run the JAGS model with 1000 adaptation steps, 10,000 burn-in iterations, saving 1000 steps, 3 MCMC chains, and thinning each chain by a factor of 10. By not specifying the number of iterations, we run the model with the default 10,000 iterations. We will also give `run_model()` a list of other parameters to track. Here, we will track the parameters `n, n3 taunoise, strata, B.X`, and `beta.X.` By default, run_model() will always track the parameter n, no matter what other parameters are specified (even if n is not explicitly specified), as this parameter is needed to calculate annual indices and trends later in the analysis. In this case, we also track n3, which we could later used to decompose the trajectory into both the GAM smooth trajectory and GAM smooth with yearly fluctuations.

~~~
# Run JAGS
jags_mod <- run_model(jags_data = jags_data,
                      n_adapt = 1000,
                      n_saved_steps = 1000,
                      n_burnin = 10000,
                      n_chains = 3,
                      n_thin = 10,
                      parallel = FALSE,
                      parameters_to_save = c(“n”,
                                             “n3”,
                                             “taunoise”,
                                             “strata”,
                                             “B.X”,
                                             “beta.X”))
~~~

Depending on what species is being modelled and what model is being used, this step in the analysis process can take up to a few days to run. If multiple processor cores are available on the user’s computer, this processing time can be reduced with the argument `parallel = TRUE`, where each MCMC chain will be run on a separate core. In any case, if the user’s geographical region permits it, the authors recommend the user pack up some granola bars, go outside, and search (and listen) for Wood Thrush in their personal birding patch while the model runs.

#### 1.3.2 Model summary and Visualizations

When the JAGS model is complete, the `run_model()` function will send a warning if any of the tracked parameters have an 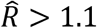. When the authors ran this worked example, convergence warnings were issued for only one n value and two n3 values; the remaining convergence warnings were for other parameters. As mentioned before, if all or most of the n values are converged, then that is sufficient for generating indices and trend estimates. In any case, if the user were wanting to run the model for more iterations in an attempt to improve convergence, they can make use of the `extract_final_values(jags_mod = jags_mod)` function and use these values as initial values for a new JAGS model.

Once the user is happy with convergence, indices and trends can be calculated. For this example, our goal is to use *bbsBayes* to create 4 figures: 1) a plot of the continental annual indices of abundance with observed means for Wood Thrush between 1966 and 2018, 2) a 2-panel plot of the national annual indices for Canada and US, 3) a geofacet plot, and 4) a heat map of population trends for Wood Thrush from 2008 - 2018 (i.e., a 10-year trend) for each stratum. We would also like to make a statement regarding the probability that the species has declined across its range by 0% (i.e., any decrease at all), by 50%, and by 100% from 2008 – 2018.

Let us begin by generating the indices of abundance for the continental, national, and stratum levels.

~~~
# Generate indices at the continental, national, and stratum
level
~~~

~~~
indices <- generate_indices(jags_mod = jags_mod,
                            jags_data = jags_data,
                            regions = c(“continental”,
                                        “national”,
                                        “stratum”))
~~~

Already, we can generate the first two figures that we specified above. Let us first create the list of trajectory plots with observed means. We will also specify some different sizing for titles and axes:

~~~
# Create a list of trajectory plots, with observed means
plot_list <- plot_indices(indices_list = indices,
                          species = “Wood Thrush”,
                          add_observed_means = TRUE,
                          title_size = 18,
                          axis_title_size = 14,
                          axis_text_size = 12)
~~~

We can then access the continental plot list with `plot_list$Continental`. The resulting plot is shown in Figure 2.

**Figure 2:**
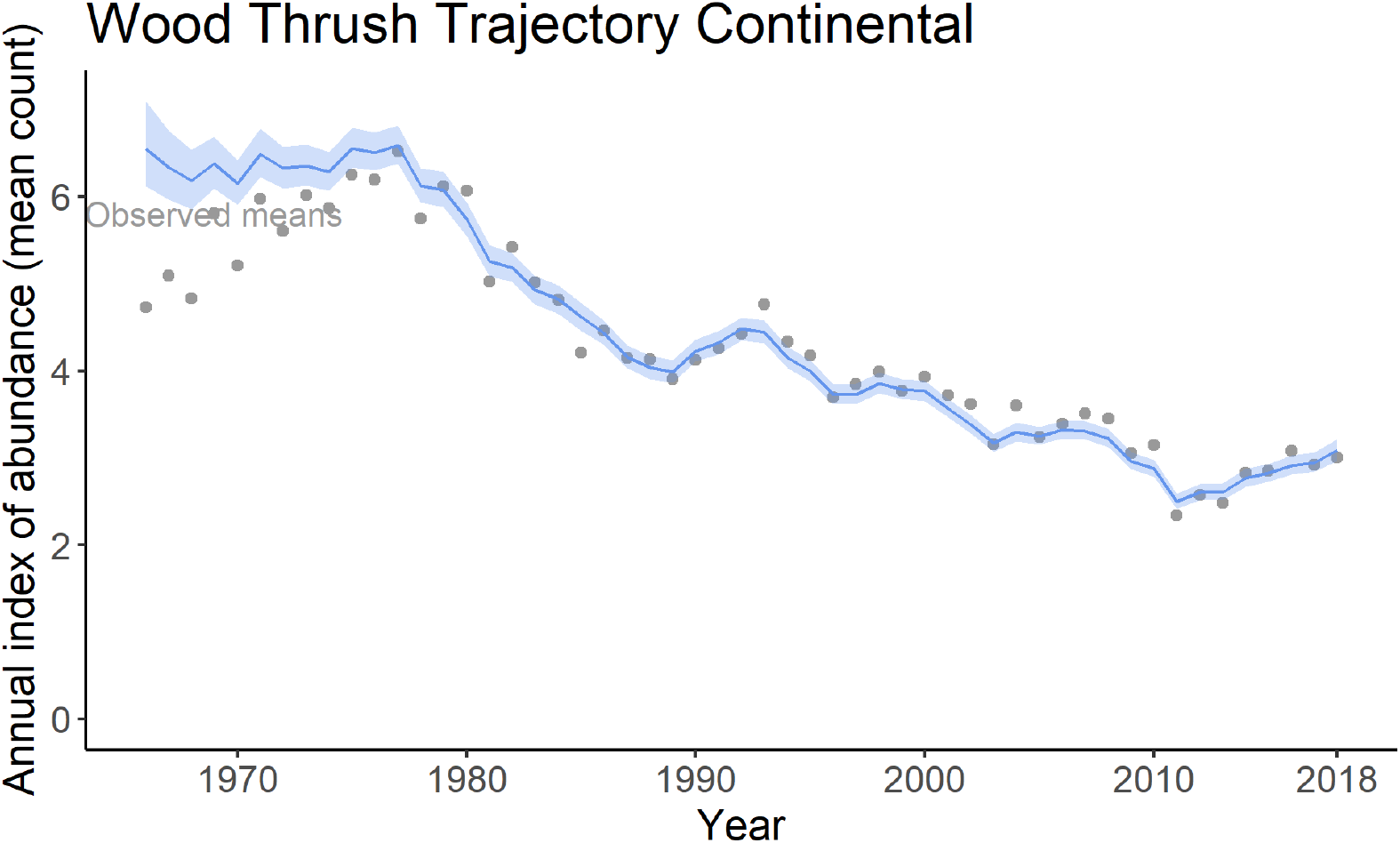
Plot of continental annual index of abundance for Wood Thrush from 1966 to 2018 with 95% credible band and observed means. This plot was generated using the `plot_indices()` function.

Generating the 2-panel plot of the national indices requires slightly more work and requires the use of the library gridExtra [1]. For the national trends, we can access the Canada and US plot with `plot_list$Canada` and `plot_list$United_States_of_America`, respectively. Thus, by using the `grid.arrange ()` function from gridExtra, we can generate a 2-panel national trajectory plot with

~~~
library(gridExtra)
grid.arrange(plot_list$Canada,
             plot_list$United_States_of_America,
             nrow = 2, ncol = 1)
~~~

which can be seen in Figure 3.

**Figure 3:**
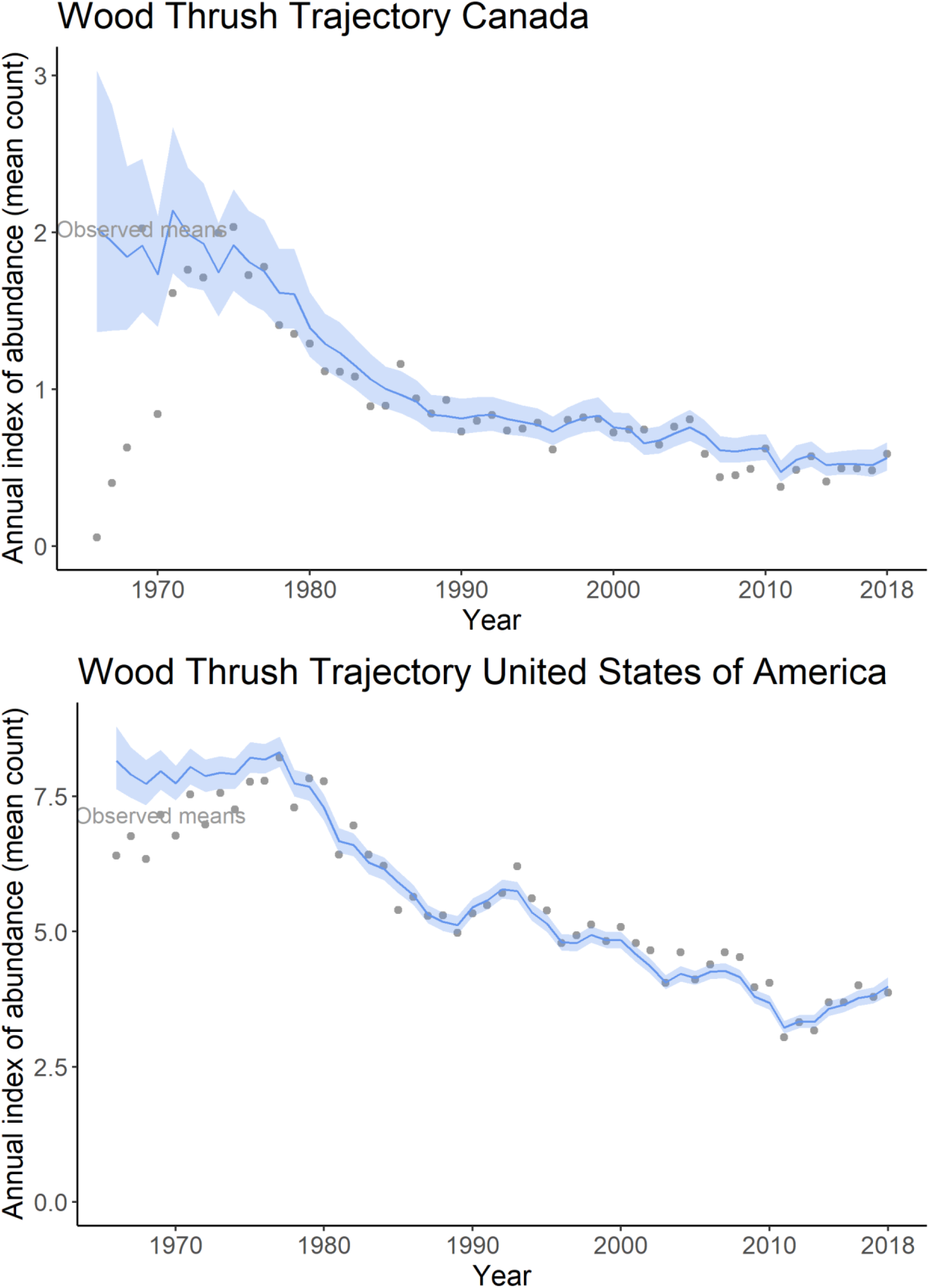
Plots of national annual indices of abundance for Wood Thrush from 1966 to 2018 for Canada and USA. These plots were generated using the `plot_indices()` function.

To create the geofacet plot given the stratification we used (recall we set `s = “bbs_cws”`), we can run

~~~
geofacet_plot(indices_list = indices,
              stratify_by = s,
              select = TRUE,
              multiple = TRUE,
              species = “Wood Thrush”)
~~~

In this case, we must set `select = TRUE` to indicate that the function must select the stratum-level data out of a data frame that contains other region types; in our case, our indices data frame contains indices at the stratum level, national level, and continental level. Additionally, we must set `multiple = TRUE` to indicate that each province/state facet may be made up of multiple strata-level trajectories that must be combined. The resulting figure can be seen in Figure 4.

**Figure 4:**
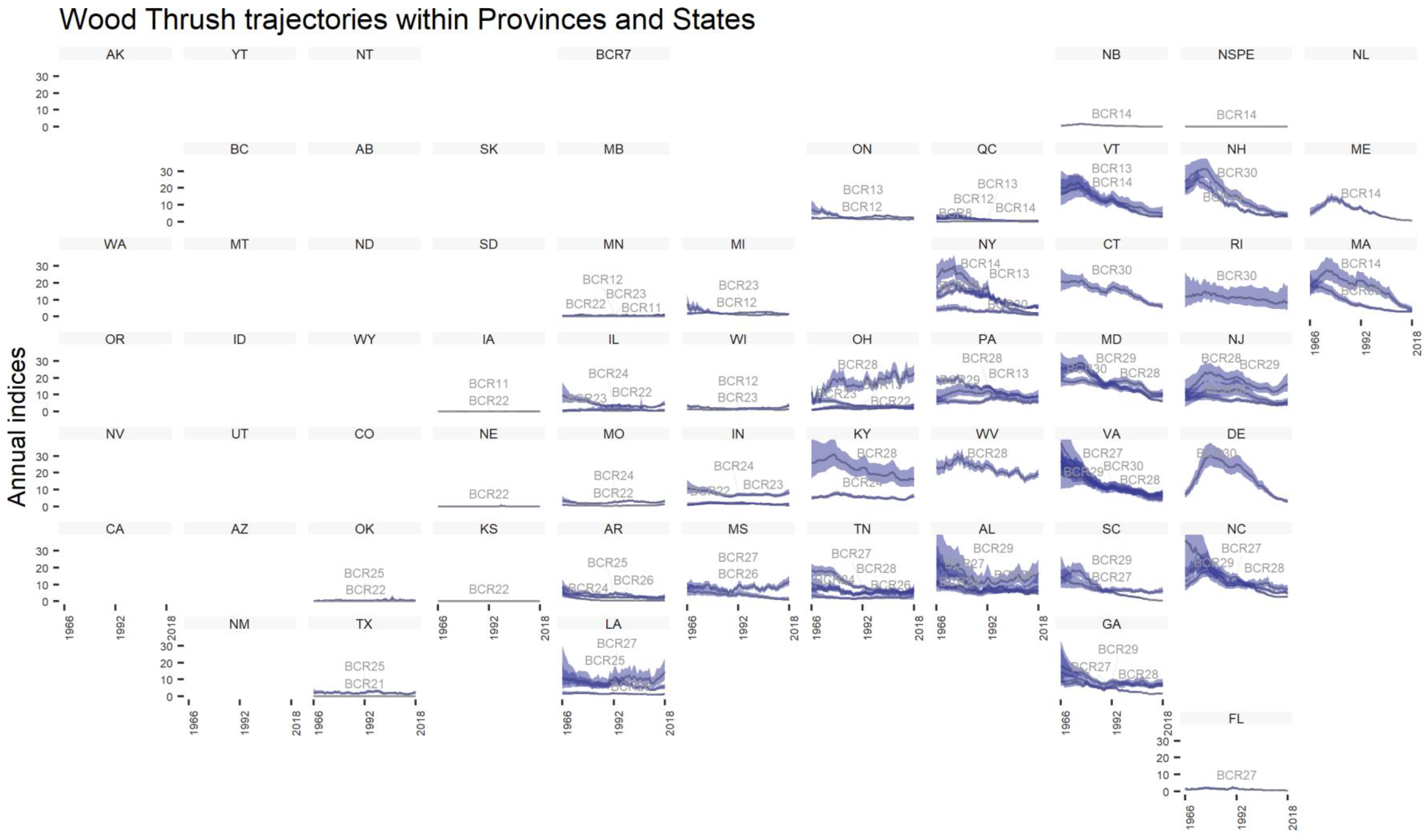
Geofacet plot of Wood Thrush trajectories for each province, state, and territory, created using the `plot_geofacet()` function.

Shifting gears slightly, let us now do some analysis of trends. As mentioned, we want to create a heat map of the trend by each stratum, and make some quantitative statements about the trends. First, we will generate the trends from 2009 to 2018

~~~
trends <- generate_trends(indices = indices,
                          Min_year = 2008,
                          Max_year = 2018,
                          prob_decrease = c(0, 50, 100))
~~~

Right away, we can generate a heat map of the trends for each stratum over this 10-year period with

~~~
generate_map(trend = trends,
                         select = TRUE,
                         stratify_by = s,
                         species = “Wood Thrush”)
~~~

Similar to the geofacet plot, we must set select = TRUE to specify that we are providing a data frame that contains trends for more than just the stratum regions, and we must also set `stratify_by = s` to indicate the original stratification used. The resulting figure is seen in Figure 5.

**Figure 5:**
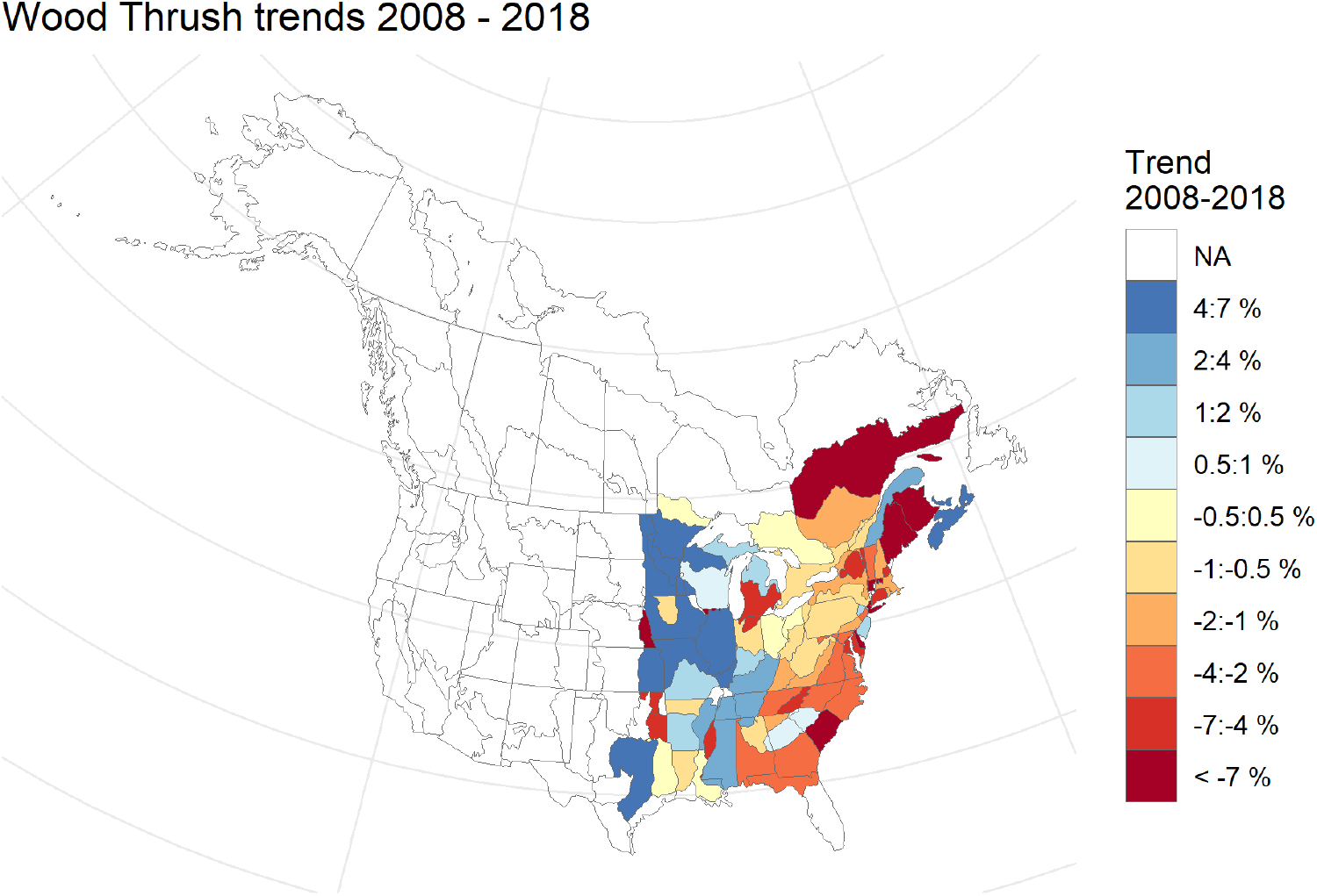
Heat map of Wood Thrush trends for each stratum for the 10-year period between 2009 and 2018. This map was generated using the `generate_map()` function.

When we generated the trends, we have also specified `prob_decrease = c(0, 50, 100)`; our trends data frame will therefore contain the posterior probabilities that Wood

Thrush has decreased by at least 0%, by at least 50%, and by at least 100% for this 10 year period for each region initially specified. Table 2 summarizes these percent changes and probabilities of change for the continent, Canada, USA, and four select strata. With these data, we can make a statement such as “our model shows a 96% probability of any population decrease at the continental level”, or “out model shows a 2% probability of the population decreasing by 50% in the CA-QC-12 stratum”.

**Table 2:**
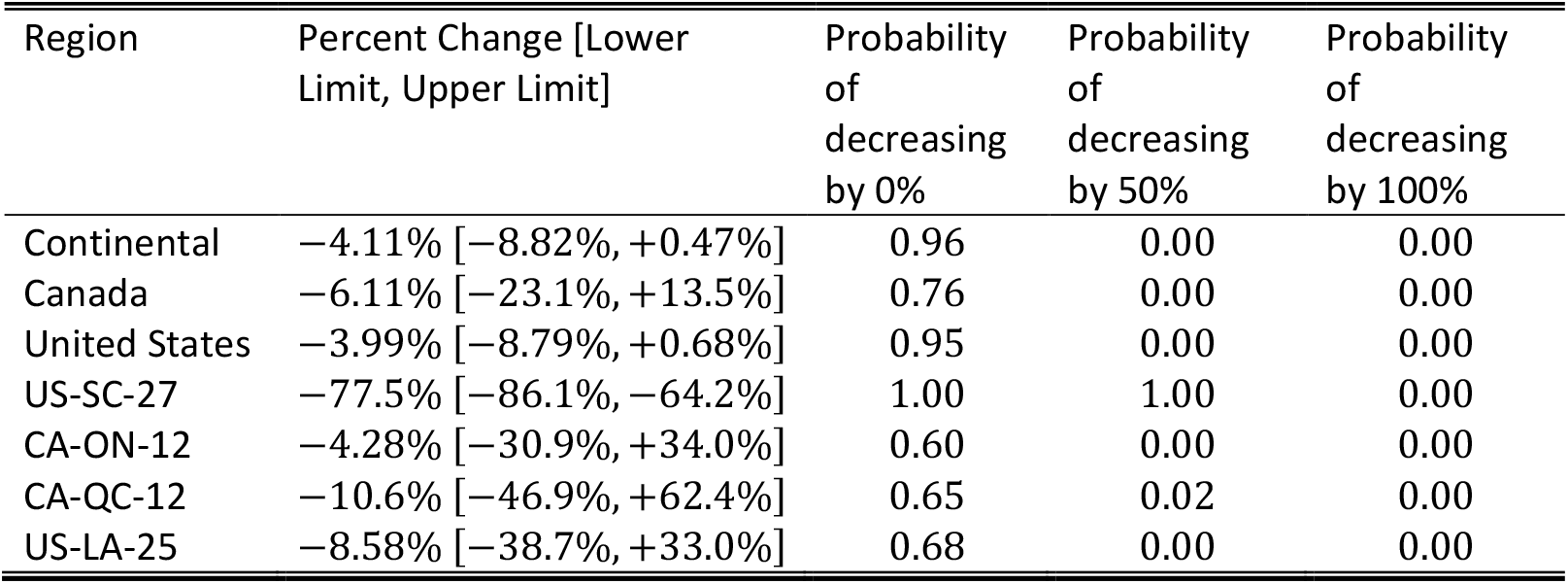
Percent change (and 95% credible interval) and probability of changes for the continent-wide trend, national trends, and stratum-level trends for select strata for Wood Thrush between 2008 – 2018. Based on the function that was run, these probabilities show the probability of the Wood Thrush population decreasing by 0%, 50%, and 100% in each of the geographical regions.

| Region | Percent Change [Lower Limit, Upper Limit] | Probability of decreasing by 0% | Probability of decreasing by 50% | Probability of decreasing by 100% |
| --- | --- | --- | --- | --- |
| Continental | −4.11% [−8.82%, +0.47%] | 0.96 | 0.00 | 0.00 |
| Canada | −6.11% [−23.1%, +13.5%] | 0.76 | 0.00 | 0.00 |
| United States | −3.99% [−8.79%, +0.68%] | 0.95 | 0.00 | 0.00 |
| US-SC-27 | −77.5% [−86.1%, −64.2%] | 1.00 | 1.00 | 0.00 |
| CA-ON-12 | −4.28% [−30.9%, +34.0%] | 0.60 | 0.00 | 0.00 |
| CA-QC-12 | −10.6% [−46.9%, +62.4%] | 0.65 | 0.02 | 0.00 |
| US-LA-25 | −8.58% [−38.7%, +33.0%] | 0.68 | 0.00 | 0.00 |

### 1.4 Quality control

The original code of *bbsBayes*, prior to being used in this package, has been continuously developed and used for several years to provide yearly trend estimates of BBS data [26, 27]. *bbsBayes* also underwent 6 months of beta testing where users could submit bugs or suggestions through GitHub issues.

The R package *testthat* [29] was used in a test harness for data fetching. Finally, 1.3 provides a worked example that a user can work through to ensure the package is functioning correctly.

A small amount of sample data is provided for the users to test the toy examples given in the documentation.

The authors of the package will continue to review bug fixes and suggested changes made in pull requests to the Github repository.

## (2) Availability

### 2.1 Operating system

Any system that can run R (obtained from https://www.r-project.org/) and JAGS (obtained from http://mcmc-jags.sourceforge.net/).

### 2.2 Programming language

R version 2.10 or higher. JAGS version 4.3.0

### 2.3 Additional system requirements

An internet connection is required to install the *bbsBayes* package, to install JAGS, and to download the BBS data. Since the BBS data is extremely data rich, it is recommended to have 8 GB or more of RAM to handle the large process created by JAGS for a given species.

### 2.4 Dependencies

R package: *progress* [5]*, jagsUI* [11]*, ggrepel* [25]*, geofacet* [9]*, ggplot2* [30]*, stringr* [31]*, grDevices* [18]*, rgdal* [3]*, dplyr* [32]*, sf* [16]*, tools* [18]*, latticeExtra* [21]*, rappdirs* [19]

### 2.5 List of contributors

This package was created by Adam C. Smith and Brandon P.M. Edwards.

### 2.6 Software location

#### 2.6.1 Archive

(e.g. institutional repository, general repository) (required – please see instructions on journal website for depositing archive copy of software in a suitable repository)

***Name:*** CRAN
***Persistent identifier:*** https://cran.r-project.org/web/packages/bbsBayes/index.html
***Licence:*** MIT
***Publisher:*** Brandon P.M. Edwards
***Version published:***
***Date published:*** 27/05/2020

#### 2.6.2 Code repository

(e.g. SourceForge, GitHub etc.) (required)

***Name:*** GitHub
***Identifier:*** https://github.com/BrandonEdwards/bbsBayes
***Licence:*** MIT
***Date published:*** 31/03/20

### 2.7 Language

R [18], JAGS [17]

## (3) Reuse potential

The goal of *bbsBayes* is to make the analysis of BBS data using hierarchical Bayesian models more accessible to the conservation community, allowing users who are familiar with R to generate population trends and trajectories for any of the species monitored by the BBS. Because the BBS represents the best population monitoring information for most species of birds in North America [6, 24], *bbsBayes* will have reuse potential in many settings, including academic, government, and conservation NGOs. This package allows an easy-to-use function to download route-level for researchers to use in custom analyses, or to run the models provided by the package. This package also gives the user the capability to download stop-level data for researchers requiring that type of data. However, *bbsBayes* does not support any analysis for stop-level data; it is for convenience only.

For users who are already familiar with analysing BBS data, *bbsBayes* will streamline the process of fetching updated BBS data, and running additional models. For users of the trend and trajectory estimates published by the CWS and USGS, *bbsBayes* makes accessible the models and data-preparation steps underlying the estimates. Finally, for researchers, the customization options that *bbsBayes* includes will be extremely useful for modelling the almost infinite set of ecological mechanisms and hypotheses that could be explored with one of the greatest long-running, rigorously-collected, continental-scale, ecological monitoring datasets in the world.

## Acknowledgements

We thank the thousands of skilled volunteers who have contributed to the Breeding Bird Survey over the years, as well as those who have served as provincial and territorial coordinators. Thank you to Dr Gregory M. Mitchell and Dr Scott Wilson (Environment and Climate Change Canada) for their comments and suggestions in the initial development phase. Thank you to Dr Maxwell B. Joseph (Earth Lab) for contributions made to data fetching functionality. We acknowledge that the National Wildlife Research Centre, on the Carleton University campus, resides on the traditional and unceded territory of the Algonquin nation.

## Copyright Notice

Authors who publish with this journal agree to the following terms:

Authors retain copyright and grant the journal right of first publication with the work simultaneously licensed under a Creative Commons Attribution License that allows others to share the work with an acknowledgement of the work’s authorship and initial publication in this journal.

Authors are able to enter into separate, additional contractual arrangements for the non-exclusive distribution of the journal’s published version of the work (e.g., post it to an institutional repository or publish it in a book), with an acknowledgement of its initial publication in this journal.

By submitting this paper you agree to the terms of this Copyright Notice, which will apply to this submission if and when it is published by this journal.

## References

[1] Auguie B. (2017). gridExtra: Miscellaneous Functions for “Grid” Graphics. R package version 2.3. https://CRAN.R-project.org/package=gridExtra

[2] Bird Studies Canada and North American Bird Conservation Initiative. (2014). Bird Conservation Regions. Published by Bird Studies Canada on behalf of the North American Bird Conservation Initiative. http://www.birdscanada.org/research/gislab/index.jsp?targetpg=bcr Accessed: 4 January 2019

[3] Bivand, R., Keitt, T., Rowlingson, B. (2018). rgdal: Bindings for the ‘Geospatial’ Data Abstraction Library. R package version 1.3-6. https://CRAN.R-project.org/package=rgdal

[4] Crainiceanu CM, Ruppert D, Wand MP. (2005). Bayesian Analysis for Penalized Spline Regression Using WinBUGS. Journal of Statistical Software, 14 (14). doi:10.18637/jss.v014.i14

[5] Csárdi, G., FitzJohn, R. (2018). progress: Terminal Progress Bars. R package version 1.2.0. https://CRAN.R-project.org/package=progress

[6] Downes, C. M., Hudson, M.A.R., Smith, A.C., and Francis, C.M. (2016). The Breeding Bird Survey at 50: scientists and birders working together for bird conservation. Avian Conservation and Ecology 11(1):8. doi: 10.5751/ACE-00855-110108

[7] Evans, M., E. Gow, R. R. Roth, M. S. Johnson, and T. J. Underwood (2011). Wood Thrush (Hylocichla mustelina), version 2.0. In The Birds of North America (A. F. Poole, Editor). Cornell Lab of Ornithology, Ithaca, NY, USA. doi: 10.2173/bna.246

[8] Gelman, A and D.B. Rubin. (1992). Inference from Iterative Simulation Using Multiple Sequences. Statistical Science 7(4), pp.457–472. doi: 10.1214/ss/1177011136

[9] Hafen, R. (2019). geofacet: ‘ggplot2’ Faceting Utilities for Geographical Data. R package version 0.1.10. https://CRAN.R-project.org/package=geofacet

[10] Hudson, M.A.R., Francis, C.M., Campbell, K.J., Downes, C.M., Smith, A.C., Pardieck, K.L. (2017). The role of the North American Breeding Bird Survey in conservation. The Condor, 119(3), pp.526–545. doi:doi.org/ 10.1650/CONDOR-17-62.1

[11] Kellner, K. (2018). jagsUI: A Wrapper Around ‘rjags’ to Streamline ‘JAGS’ Analyses. R package version 1.5.0. https://CRAN.R-project.org/package=jagsUI

[12] Knape, J. (2016). Decomposing trends in Swedish bird populations using generalized additive mixed models. Journal of Applied Ecology, 53, pp.1852–1861. doi: 10.1111/1365-2664.12720

[13] North American Bird Conservation Initiative (2016). The State of North America’s Birds 2016. Ottawa: Environment and Climate Change Canada, pp.1–8.

[14] North American Bird Conservation Initiative Canada (2019). The State of Canada’s Birds 2019. Ottawa: Environment and Climate Change Canada, pp.1–12.

[15] Pardieck, K.L., D.J. Ziolkowski Jr., M. Lutmerding, V. Aponte and M-A.R. Hudson. (2019). North American Breeding Bird Survey Dataset 1966 - 2018, version 2018.0. U.S. Geological Survey, Patuxent Wildlife Research Center. doi: 10.5066/P9HE8XYJ

[16] Pebesma, E. (2018). Simple Features for R: Standardized Support for Spatial Vector Data. The R Journal 10(1), pp.439–446. doi: 10.32614/RJ-2018-009

[17] Plummer, MA. (2003). JAGS: A program for analysis of Bayesian graphical models using Gibbs sampling.

[18] R Core Team (2018). R: A language and environment for statistical computing. R Foundation for Statistical Computing, Vienna, Austria. URL https://www.R-project.org/.

[19] Ratnakumar, S., Mick, T., Davis, T. (2016). rappdirs: Application Directories: Determine Where to Save Data, Caches, and Logs. R package version 0.3.1. https://CRAN.R-project.org/package=rappdirs

[20] Rosenberg, K.V., Dokter, A.M., Blancher, P.J., Sauer, J.R., Smith, A.C., Smith, P.A., Stanton, J.C., Panjabi, A., Helft, L., Parr, M., Marra, P.P. (2019). Decline of the North American avifauna. Science 366(6461), pp.120–124. doi: 10.1126/science.aaw1313

[21] Sarkar, D and Andrews, F (2016). latticeExtra: Extra Graphical Utilities Based on Lattice. R package version 0.6-28. https://CRAN.R-project.org/package=latticeExtra

[22] Sauer, J.R. and Link, W.A., (2003). Hierarchical models and the analysis of bird survey information. Ornis Hungarica, 12-13(1-2), pp.217–222.

[23] Sauer, J.R. and Link, W.A., (2011). Analysis of the North American Breeding Bird Survey using hierarchical models. Auk, 128(1), pp.87–98. doi: 10.1525/auk.2010.09220

[24] Sauer, J.R., Pardieck, K.L., Ziolkowski, D.J., Smith, A.C., Hudson, M.A.R., Rodriguez, V., Berlanga, H., Niven, D.L., Link, W.A. (2017). The first 50 years of the North American Breeding Bird Survey. The Condor, 119(3), pp.576–593. doi: 10.1650/CONDOR-17-83.1

[25] Slowikowski, K. (2020). ggrepel: Automatically Position Non-Overlapping Text Labels with ‘ggplot2’. R package version 0.8.2. https://CRAN.R-project.org/package=ggrepel

[26] Smith, A.C., and Edwards, B.P.M. (2020). Improved status and trend estimates from the North American Breeding Bird Survey using a hierarchical Bayesian generalized additive model. bioRxiv 2020.03.26.010215. doi: 10.1101/2020.03.26.010215

[27] Smith, A.C., Hudson, M.A.R., Downes, C., and Francis C.M. (2014). Estimating breeding bird survey trends and annual indices for Canada: how do the new hierarchical Bayesian estimates differ from previous estimates? The Canadian Field-Naturalist, 128(2), pp.119–134. doi: 10.22621/cfn.v128i2.1565

[28] United States Geological Survey, 2019. BBS Bibliography. Available at https://www.pwrc.usgs.gov/BBS/Bibliography/

[29] Wickham, H. (2011). testthat: Get Started with Testing. The R Journal, 3(1) pp.5–10

[30] Wickham, H. (2016). ggplot2: Elegant Graphics for Data Analysis. Springer-Verlag New York.

[31] Wickham, H. (2018). stringr: Simple, Consistent Wrappers for Common String Operations. R package version 1.3.1. https://CRAN.R-project.org/package=stringr

[32] Wickham, H., François, R., Henry, L., Müller, K. (2019). dplyr: A Grammar of Data Manipulation. R package version 0.8.3. https://CRAN.R-project.org/package=dplyr

[33] Link, W.A., J.R. Sauer, and D.K. Niven. (2017). Model selection for the North American Breeding Bird Survey: A comparison of methods. Condor 119(3):546–556. doi: 10.1650/CONDOR-17-1.1

